# Exercise training at different intensities induces heat stress, disrupts barrier function and alters microbiota in the gut of mice

**DOI:** 10.1101/2024.07.10.602866

**Authors:** Puqiao Lian, Artemiy Kovynev, Lei Wang, Amanda C. M. Pronk, Aswin Verhoeven, Martin Giera, Suzan Thijssen, Borja Martínez Téllez, Sander Kooijman, Patrick C. N. Rensen, Harro Timmerman, Harry J. Wichers, Paul A. J. Henricks, Gert Folkerts, Milena Schönke, Saskia Braber

## Abstract

Exercise is generally beneficial for health but strenuous exercise can have detrimental effects on the gastrointestinal tract. The combination of ischemia and heat shock during exercise is a crucial contributor to intestinal epithelial damage. Growing evidence points towards an important regulatory role of gut microbes in intestinal homeostasis. Here, we characterize and compare the effects of moderate and vigorous exercise training on intestinal epithelial damage, stress response, inflammatory response, and gut microbiota alterations in mice and investigate the mechanisms underlying exercise-induced intestinal injury. Exercise training for six weeks caused heat stress in the intestine, resulting in the disruption of the intestinal epithelial barrier and local inflammation. This was characterized by increased colonic HSP-70 and HSF-1 protein expression, increased epithelial permeability, decreased colonic expression of tight junction proteins ZO-1 and occludin and intestinal morphological changes. Daily moderate exercise training caused hereby more severe injury than vigorous training on alternating days. Furthermore, exercise training altered the gut microbiota profile. The abundance of *Lactobacillaceae* was reduced, potentially contributing to the deteriorated intestinal status, while the abundance of short-chain fatty acid-producing *Lachnospiraceae* was increased, especially following vigorous training. This increase in short-chain fatty acid-producing bacteria following vigorous training possibly counteracted the impairment of the intestinal barrier function. In summary, exercise disrupts the intestinal barrier function, with vigorous exercise training with intermittent rest days being less damaging than daily moderate exercise training.

## 1. Introduction

Exercise has favourable effects on the human body and its benefits on the musculoskeletal, cardiovascular, and endocrine systems are well-documented [1]. However, an increasing number of studies points to simultaneous adverse effects of exercise on the gastrointestinal (GI) tract and our previous study demonstrated that endurance exercise increases the gut permeability in healthy volunteers [2]. Strenuous exercise, such as marathon running, cycling or triathlon events, is considered a contributing factor to gastrointestinal symptoms and can induce the so-called exercise-induced GI syndrome [3]. This syndrome, characterized by bloating, nausea or diarrhoea, is caused by two distinct pathways: the neuroendocrine-GI pathway and the circulatory-GI pathway [4,5]. The first pathway is associated with an increase in sympathetic activation and a consequent inhibition of GI function. The core of the latter pathway is the redistribution of blood flow to working muscles and peripheral circulation, reducing visceral perfusion. Exercise can lead to a reduction in superior mesenteric artery blood flow by up to 43% in humans [6,7]. This results in local ischemia and hypoxia in the intestine, increasing the production of reactive oxygen species (ROS) and activating signalling pathways that lead to increased gut permeability [8]. Moreover, exercise raises the body core temperature and causes heat stress that activates signalling pathways that reduce epithelial barrier function and causes widespread damage to intestinal epithelium, including shrinking and sloughing of villi [9,10]. Although there is great inter-individual variation, the threshold and degree of intestinal injury may correlate with exercise intensity and environmental temperature [11,12].

The negative impact of strenuous exercise on the GI environment involves disruption of the intestinal epithelial barrier integrity formed by protein complexes called tight junctions (TJs) and adherens junctions (AJs). These junctional complexes form a “seal” between adjacent cells and act as gatekeeper of the gut barrier. Our previous study showed that combined hypoxic and hyperthermal exposure to co-cultured colonic epithelial cells increases paracellular permeability and activates oxidative stress and heat stress pathways [13]. Heat shock proteins (HSPs), acting as molecular “chaperones” protecting cells from environmental stress, are upregulated under heat stress, assisting with protein synthesis, assembly, and degradation and maintaining the viability and proliferative capacity of cells [14]. Increased HSPs observed in the gut of athletes after endurance competitions prevent TJ breakdown and protect the cytoskeleton of intestinal epithelial cells from hypoxia and hyperthermia-related damage [15,16].

In addition, exercise training modulates the composition of the intestinal microbiota which may affect the normal intestinal function [17,18], however, the underlying mechanisms of this effect remain elusive. This is the first murine study that compares intestinal alterations induced by exercise training at different intensities (moderate and vigorous) and durations (2, 4 and 6 weeks) and investigates the mechanisms behind exercise-induced intestinal injury. The aim was to identify the key regulators of this “destruction and reconstruction process” following exercise.

## 2. Materials and Methods

### 2.1. Animals and experimental design

Male C57BL/6J mice at 9 weeks of age were randomly divided into the following groups: control (CON, n = 60), moderate treadmill exercise training (MOD-EX, n = 60), and vigorous treadmill exercise training (VIG-EX, n = 60). In each group, the mice were randomly divided into 3 subgroups by their euthanasia time points: 2, 4, and 6 weeks of intervention (n = 20, respectively). For details see Supplementary Information.

### 2.2. Treadmill running exercise protocol

A 3-day progressive exercise training regime was adapted to familiarize the MOD-EX and VIG-EX mice to a rodent treadmill where the speed and incline of the track were incrementally increased. From day 4 to 15, the mice followed a fixed training regime. To ensure that the exercise straining remained strenuous, starting from day 15 and day 29, the running speed of the MOD-EX group and the VIG-EX group was increased. The outline of the time points for exercise training, rest and sacrifice are depicted in Supplementary Fig. S1. For details see Supplementary Information.

### 2.3. Euthanasia and intestinal fluorescein permeability

Immediately after the last treadmill training session, FITC-dextran (FD; 4 kDa) was administered to each mouse by oral gavage (500 mg/kg BW). For details see Supplementary Information.

### 2.4. Tissue collection and treatment

Two cm of the segments from the proximal, middle and distal parts of the small intestine and from the colon were isolated and cleaned with ice cold sterile PBS. The spleen was harvested and weighed. Tissue samples were snap-frozen on dry ice and stored at −80°C until future use. Swiss rolls of the intestinal tissue were and then embedded in paraffin blocks for histomorphological evaluation. For details see Supplementary Information.

### 2.5. Western blotting

The colonic segments were lysed and homogenized with cold Pierce RIPA buffer containing a protease inhibitor. For details see Supplementary Information.

### 2.6. Histomorphological evaluation

The Swiss rolls of proximal and distal parts of the small intestine and colonic parts were cut into 5 µm sections and stained with haematoxylin and eosin (H&E). The epithelial morphology, mucosal architecture and inflammation of the intestinal samples were evaluated and scored by double-blinded technicians. For details see Supplementary Information.

### 2.7. Immunofluorescence staining

After standard procedures of gradient rehydration and blocking, the sections were incubated with primary antibody against occludin and then incubated with an Alexa-Fluor fluorescently conjugated secondary antibody. The slides were mounted and counterstained with DAPI. The cells were visualized by a confocal microscopy system. For details see Supplementary Information.

### 2.8. Caecal metabolomics measurement

The concentrations of short-chain fatty acids (SCFAs) and amino acids (AAs) in the caecal contents were quantified by nuclear magnetic resonance spectroscopy. For details see Supplementary Information.

### 2.9. DNA extraction of caecal contents, library construction, gut microbiota sequencing, and bioinformatics analysis

Caecal contents were collected from individual mice and stored at −80°C until future use. For details see Supplementary Information.

### 2.10. Statistical analysis

Statistical analyses were performed by using GraphPad Prism software or the functions of R packages. Differences between groups were determined with parametric repeated measures (RM) two-way analysis of variance (ANOVA), or ordinary one-way ANOVA with Bonferroni post-hoc test, or non-parametric Kruskal-Wallis rank-sum test, two-sided Welch’s test, or permutational multivariate analysis of variance (PERMANOVA). A Pearson rank correlation test was conducted to examine associations between the parameters tested, with coefficient |R| > 0.4 being considered correlated. Results were considered statistically significant when p < 0.05 or corrected q < 0.05.

## 3. Results

### 3.1. Exercise training increases intestinal leakage and disrupts the intestinal tight junction network in mice

Exercise-induced changes in gut permeability following 2, 4 or 6 weeks of treadmill training were evaluated by measuring the leakage of the fluorescent molecule FITC-Dextran (4 kDa) into the blood after oral administration. Neither moderate (MOD-EX) nor vigorous (VIG-EX) treadmill training affected intestinal FITC-Dextran permeability after 2 weeks of training. Four weeks of MOD-EX, however, significantly increased the serum FITC-Dextran concentration, indicating increased barrier leakage across the intestinal epithelium. After 6 weeks of training, the serum FITC-Dextran concentration was significantly elevated in both MOD-EX and VIG-EX compared to the control mice (CON) (Fig. 1A).

**Figure 1.**
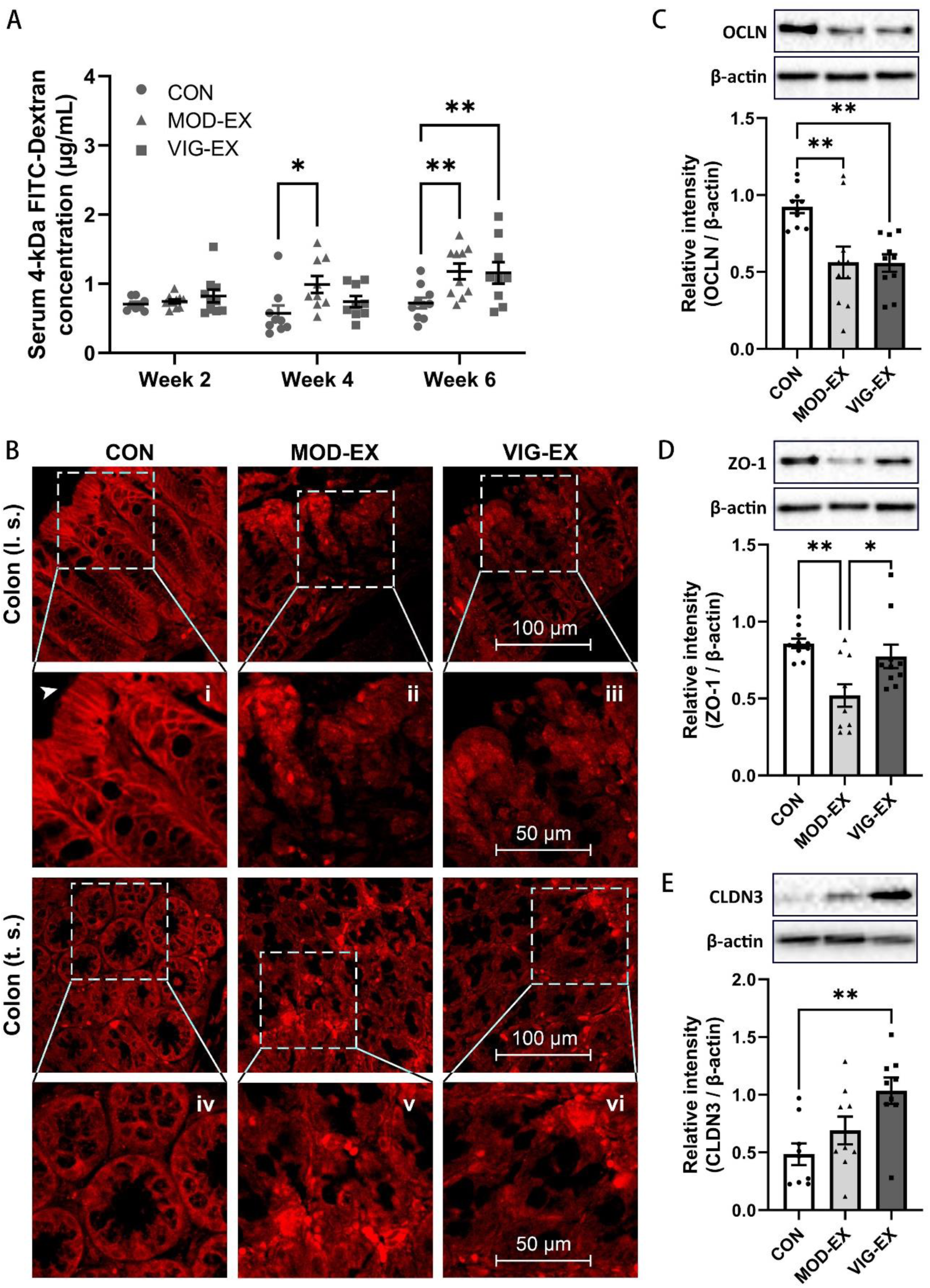
Effect of exercise on intestinal barrier function. (**A**) Serum 4-kDa FITC-Dextran concentration 1 h after oral gavage following 2, 4 and 6 weeks of exercise training. Data are presented as mean ± SEM; n = 9 (VIG-EX of Week 6) or n = 10 (other groups) per group. Statistical differences were analysed by one-way ANOVA followed by Bonferroni’s multiple comparison test. (**B**) Immunofluorescence staining of colonic occludin protein after 6 weeks of exercise training. The results were acquired by Leica TCS SP8 confocal microscope with HCX IRAPO L 25×/0.95 objective lens at 2.25×digital magnification; pinhole: 1.50 AU. Red colour: fluorescent signals from Alexa-Fluor 594 secondary antibodies. (**C**, **D**, **E**) Relative protein expression of colonic claudin-3, occludin and ZO-1 after 6 weeks of exercise training, assessed by Western blot. All target proteins were normalized to reference protein β-actin. Data are presented as mean ± SEM; n = 9-10 per group. Statistical differences were analysed by one-way ANOVA followed by Bonferroni’s multiple comparison test. * p < 0.05, ** p < 0.01. CON: Control; MOD-EX: moderate exercise; VIG-EX: vigorous exercise; l.s.: longitudinal section; t.s.: transverse section; OCLN: occludin; ZO-1: zonula occludens-1; CLDN3: claudin-3.

Immunofluorescent staining for TJ protein occludin (OCLN) was conducted to visualize its localization in the colonic villus structures. In the colon of untrained CON mice, OCLN formed a continuous mesh-like pattern in longitudinal villi (Fig. 1B-i) and a clear ring pattern in cross-sectional villi (Fig. 1B-iv). A brush-like appearance of OCLN was observed in compact columnar cells at the tips of the villi in CON (Fig. 1B-i, arrowhead). Exercise training of either intensity induced a disturbed cellular distribution of OCLN and significant loss of columnar cells in mouse colonic villi (Fig. 1B-ii, -iii). Moreover, concentrated protein clusters and a more diffuse and cytosolic distribution were observed in cross sections of the villi (Fig. 1B-v, -vi). In addition, the OCLN localization was evaluated in proximal small intestinal (duodenal) and distal small intestinal (ileal) sections where group differences were less pronounced (Supplementary Fig. S3). OCLN localized at the cell membrane showed continuous chain-like structures forming the rim of normal duodenal and ileal villi in CON, which is less clear following exercise training (Supplementary Fig S3).

The changes in the TJ network after exercise were confirmed by Western blotting. Both MOD-EX and VIG-EX decreased OCLN protein expression in the mouse colon (Fig. 1C). In addition, the protein expression of ZO-1 significantly decreased in MOD-EX, while VIG-EX did not affect ZO-1 expression but enhanced CLDN3 expression (Fig. 1D, E).

### 3.2. Exercise elevates rectal temperature and induces a heat stress response in mouse colon

The thermal effects of exercise on the mouse intestine were determined by measuring rectal temperature and the expression of heat shock-related proteins/transcription factors. During the 6 weeks of treadmill training, the rectal temperature of CON narrowly fluctuated around the baseline (37.8 ± 0.1°C, mean ± SEM, *sic passim*) within approx. 0.5°C (Fig. 2A). The rectal temperatures of MOD-EX and VIG-EX were significantly increased 15 min after exercise compared to CON and reached 39.0 ± 0.1°C and 38.7 ± 0.1°C, respectively, at week 6 (Fig. 2A). However, the rectal temperature between the two exercise groups was not significantly different.

**Figure 2.**
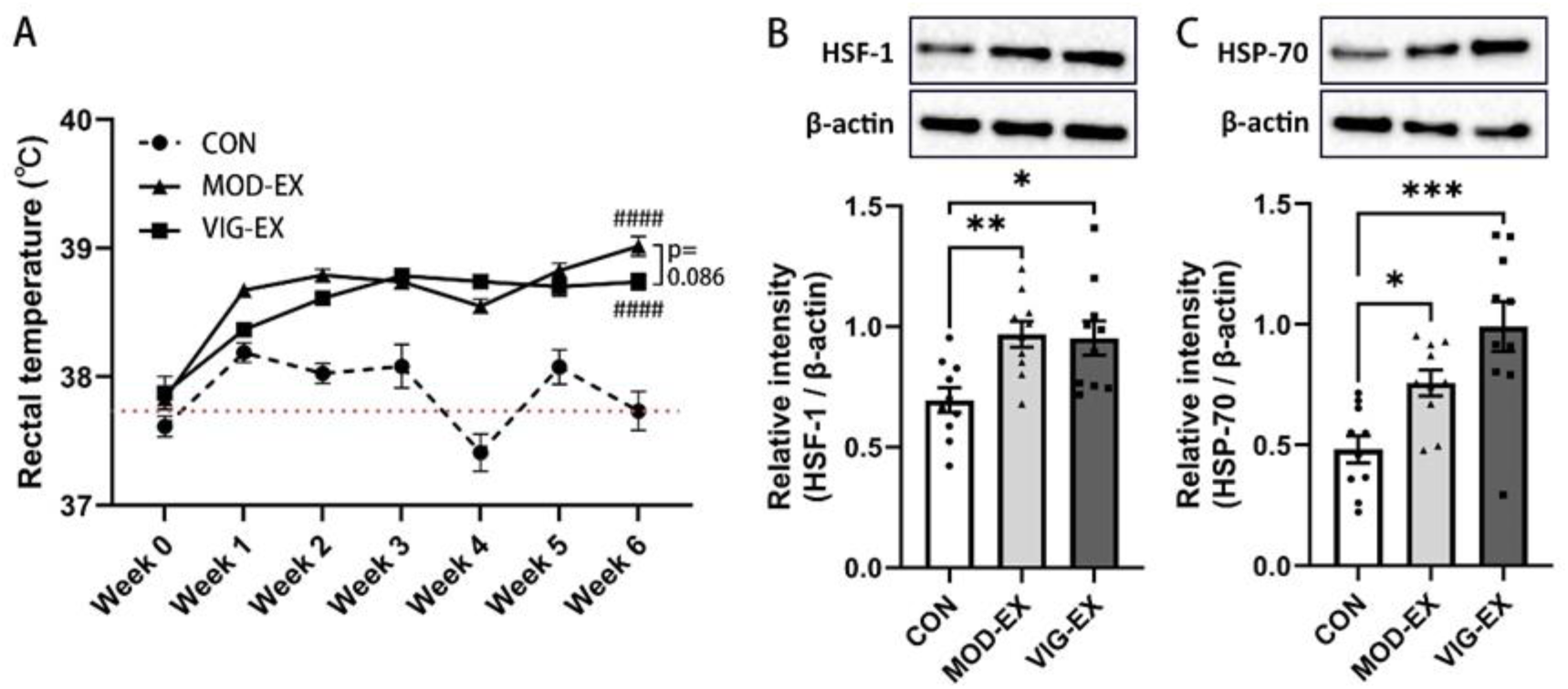
The effect of exercise on the intestinal heat stress response. (**A**) Weekly post-exercise (after 15 min) rectal temperatures of mice acquired by a rectal thermometer. The dotted red line represents the average rectal temperature of all groups. Data are presented as mean ± SEM; n = 20 per group. Statistical differences were analysed by repeated measures two-way ANOVA. #### p < 0.0001 vs Control. Relative protein expression of (**B**) HSF-1 and (**C**) HSP70 in the colon after 6 weeks of exercise training assessed by Western blot. The target proteins were normalized to reference protein β-actin. Data are presented as mean ± SEM; n = 10 per group. Statistical differences were analysed by one-way ANOVA followed by Bonferroni’s multiple comparison test. * p < 0.05, ** p < 0.01, *** p < 0.001. CON: Control; MOD-EX: moderate exercise; VIG-EX: vigorous exercise; HSP: heat shock protein; HSF: heat shock factor.

At week 6, after the final running session, the mouse colon tissue was collected and the protein levels of HSP-70, the conserved ubiquitously expressed protein of the heat shock protein family, and the transcription factor HSF-1, the primary mediator of transcriptional responses to heat stress, were determined. Both MOD-EX and VIG-EX significantly enhanced the abundance of HSF-1 in the colon (Fig. 2B). Correspondingly, the protein abundance of chaperone HSP-70 was increased in both exercise groups, without significant differences between MOD-EX and VIG-EX (Fig. 2C).

### 3.3. Exercise training alters intestinal morphology and induces local intestinal inflammation in mice

Following 6 weeks of training, proximal and distal small intestine and colonic sections were examined to determine the effect of exercise training on intestinal histomorphology. In CON, limited inflammatory infiltration was observed. CON showed well-distended villi from base to tip in the proximal small intestine (duodenum), distal small intestine (ileum) and colon, and the mucosal and submucosal architecture was intact (Fig. 3A, B). Intestinal villus length is correlated to the digestive and absorptive functions of the intestine, as it affects surface area of the digestive tract [19]. The duodenal villus length was significantly shorter in MOD-EX compared to CON (Fig. 3C, D), while this effect was less pronounced in VIG-EX. No significant changes in villus length were observed in the ileum after exercise training (Fig. 3D), but blunted and deformed villi were distinct characteristics in the duodenum and ileum (Fig. 3A, asterisk) of both exercise groups.

**Figure 3.**
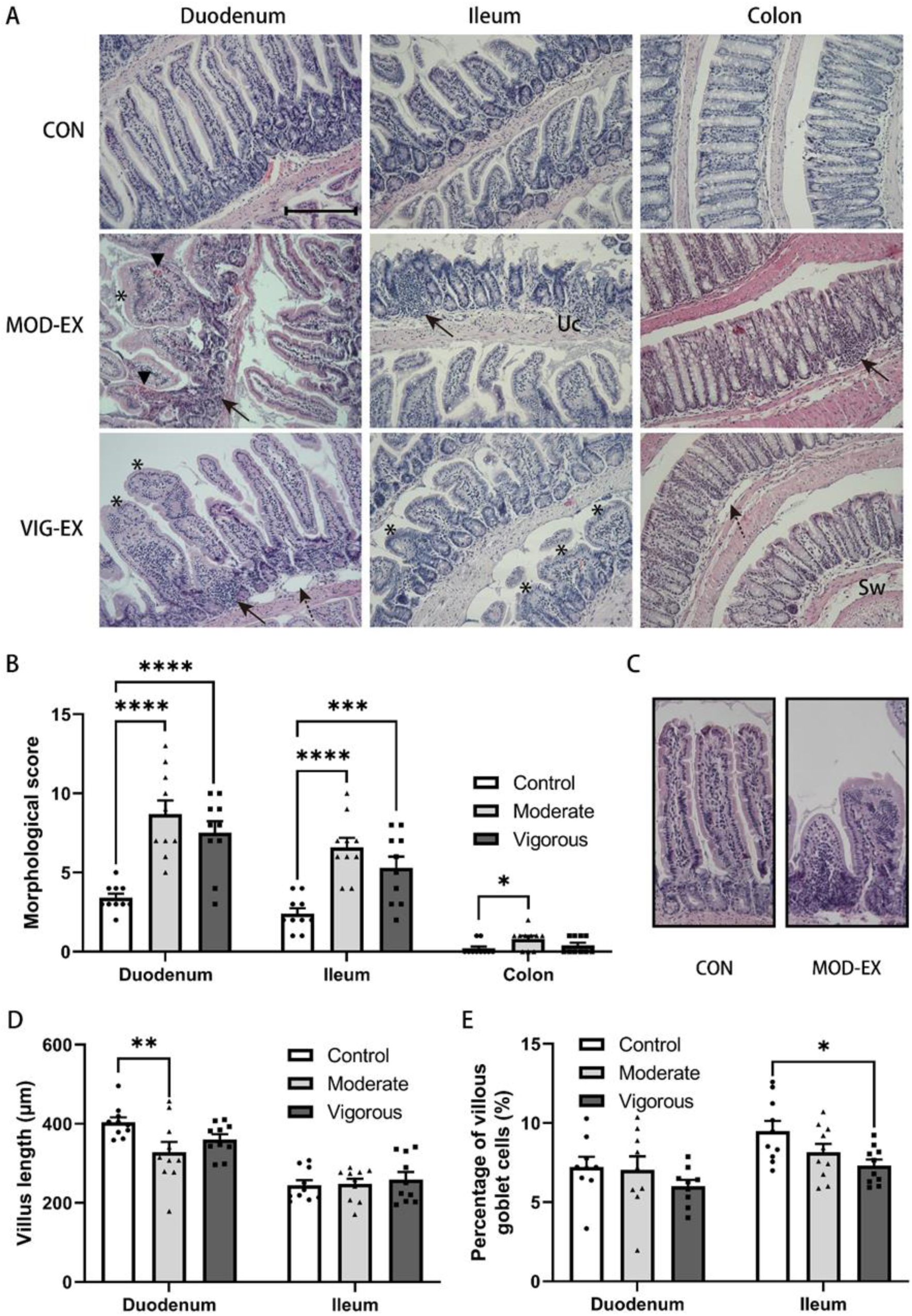
Effect of exercise training on intestinal morphology and inflammation. (**A**) Representative images of H&E-stained proximal small intestinal (duodenal), distal small intestinal (ileal), and colonic sections of mice after 6 weeks of exercise training illustrate the inflammatory cell infiltration, epithelial changes and altered mucosa architecture. The images were acquired by Olympus BX50 microscope with UPlanFI 20× /0.25 objective lens and Leica DFC320 10× camera (200×, scale bar 200 μm). Solid arrow: mucosal or submucosal infiltration of leukocytes; dashed arrow: submucosal oedema; arrowhead: haemorrhage; Uc: ulceration; asterisk: blunted villus; Sw: submucosal widening. (**B**) Histological scores of intestinal morphology based on inflammatory cell infiltration, epithelial changes and altered mucosa architecture. (**C**) Representative images of cross-sectioned duodenal villi (**D**) Villus lengths measured by ImageJ software. (**E**) Percentages of villus goblet cells counted after Alcian Blue/Nuclear Fast Red staining. Data are presented as mean ± SEM; n = 10 mice per group. Statistical differences were analysed by one-way ANOVA followed by the Bonferroni’s multiple comparison test. ** p < 0.01, *** p < 0.001, **** p < 0.0001. CON: Control; MOD-EX: moderate exercise; VIG-EX: vigorous exercise.

However, significantly fewer goblet cells were counted in the ileum of VIG-EX, suggesting a possibly impaired mucus production, while no significant differences were observed between MOD-EX and CON (Fig. 3E). No significant changes in the number of goblet cells were observed in the duodenum after exercise training (Fig. 3E).

In MOD-EX and VIG-EX, mucosal or submucosal infiltration of leukocytes combined with occasional intestinal haemorrhage was detected in the duodenum, ileum and colon (Fig. 3A, solid arrows, and arrowheads). Additionally, in MOD-EX, ulceration and submucosal widening were observed in all parts of the intestine of MOD-EX (Fig. 3A, Uc) and submucosal widening and submucosal oedema were observed in duodenal, ileal and colonic sections in VIG-EX (Fig. 3A, asterisks, solid arrows, dashed arrows and Sw). These alterations caused a significantly higher morphological score of duodenum and ileum in MOD-EX and VIG-EX and of colon in MOD-EX compared to CON (Fig. 3B).

Interestingly, the serum concentration of CRP, a marker for global inflammation secreted by the liver, was increased following 6 weeks of MOD-EX (Supplementary Fig. S4A) but not VIG-EX, suggesting mildly elevated systemic inflammation with MOD-EX. However, quantification of several circulating cytokines/chemokines showed that the levels of pro-inflammatory G-CSF, CXCL1 and CCL2 were not significantly different between the groups (Supplementary Fig. S4B-D). Spleen/body weight ratios were also not different between the groups (Supplementary Fig. S5C).

### 3.4. Vigorous but not moderate exercise training affects caecal metabolite levels

Following 6 weeks of exercise training, caecal contents were collected and the concentrations of the six most abundant SCFAs (formate, acetate, propionate, butyrate, isobutyrate and valerate), lactate and four amino acids that are universally utilized by gut microbes to produce SCFAs (alanine, methionine, valine and threonine) were determined 1 h after the final exercise bout. VIG-EX, but not MOD-EX, increased caecal levels of acetate, butyrate and propionate (Fig. 4B-D), but decreased formate and valerate concentrations (Fig. 4A, F). VIG-EX for 6 weeks decreased the levels of amino acids in the cecum, including alanine, methionine, valine and threonine (Fig. 4H-K). Accordingly, total concentration of SCFAs and AAs in VIG-EX was significantly changed compared to CON and MOD-EX (Fig. 4L, M). Neither training program affected the caecal levels of lactate (Fig. 4G). Notably, there was no significant difference in the average daily food intake and cumulative food intake over 6 weeks between the groups (Supplementary Fig. S6).

**Figure 4.**
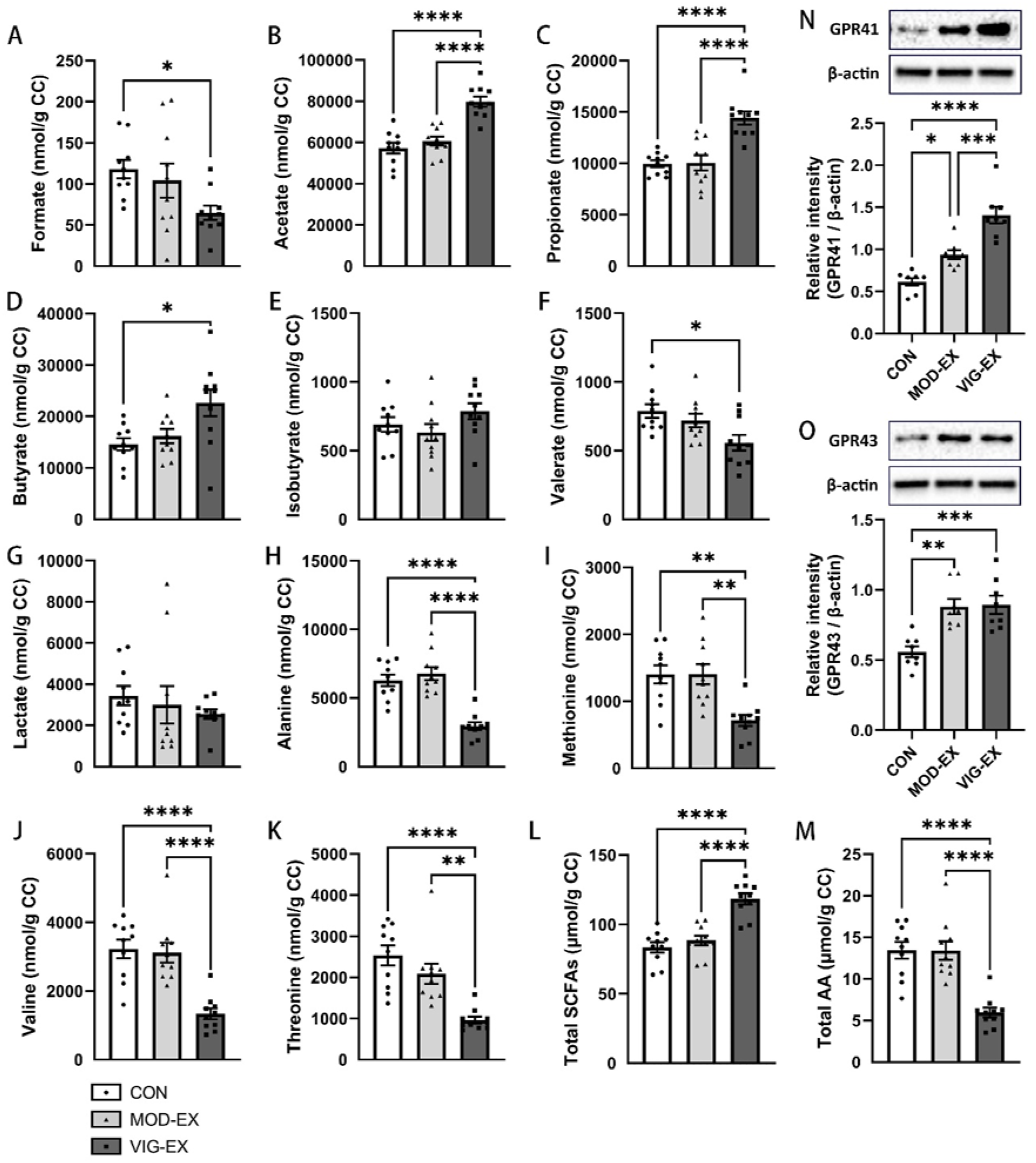
The effect of exercise on caecal metabolite levels. Concentrations of short-chain fatty acids (**A** - **F**), lactate (**G**), and amino acids (**H** - **K**) in caecal contents after 6 weeks of exercise training measured 1 h after the last training bout by nuclear magnetic resonance spectroscopy. (**L**, **M**) Concentrations of total short-chain fatty acids and total amino acids in caecal contents after 6 weeks of exercise training measured 1 h after the last training bout. (**N**, **O**) The effect of strenuous exercise on colonic G-protein-coupled receptors 41 and 43. Relative protein expression of GPR41 and GPR43 in the murine colon after 6 weeks of exercise assessed by Western blot. All target proteins were normalized to reference protein β-actin. Data in the bar plots are presented as mean ± SEM; n = 10 per group. Statistical differences were analysed by one-way ANOVA followed by the Bonferroni’s multiple comparison test. * p < 0.05, ** p < 0.01, *** p < 0.001, **** p < 0.0001. CC: caecal contents; CON: Control; MOD-EX: moderate exercise; VIG-EX: vigorous exercise; SCFAs: short-chain fatty acids; AAs: amino acids; GPR: G-protein-coupled receptor.

Both MOD-EX and VIG-EX significantly increased the protein expression of G-protein-coupled receptor (GPR) 41 and GPR43, the main mammalian receptors of SCFAs, in colon compared to CON (Fig. 4N, O). Moreover, VIG-EX increased the expression of GPR41 to a greater extent than MOD-EX.

### 3.5. Exercise training promotes changes in microbiota diversity

After 6 weeks of exercise training, the effect on the composition of the murine gut microbiota was evaluated. Significant differences in alpha diversity, showing the diversity within an individual sample, and beta diversity, showing the (dis-)similarities of the microbiomes of different groups, were evident in MOD-EX and VIG-EX when compared to CON (p = 0.001). However, the two exercise groups did not significantly differ from each other in alpha or beta diversity (Fig. 5A, B).

**Figure 5.**
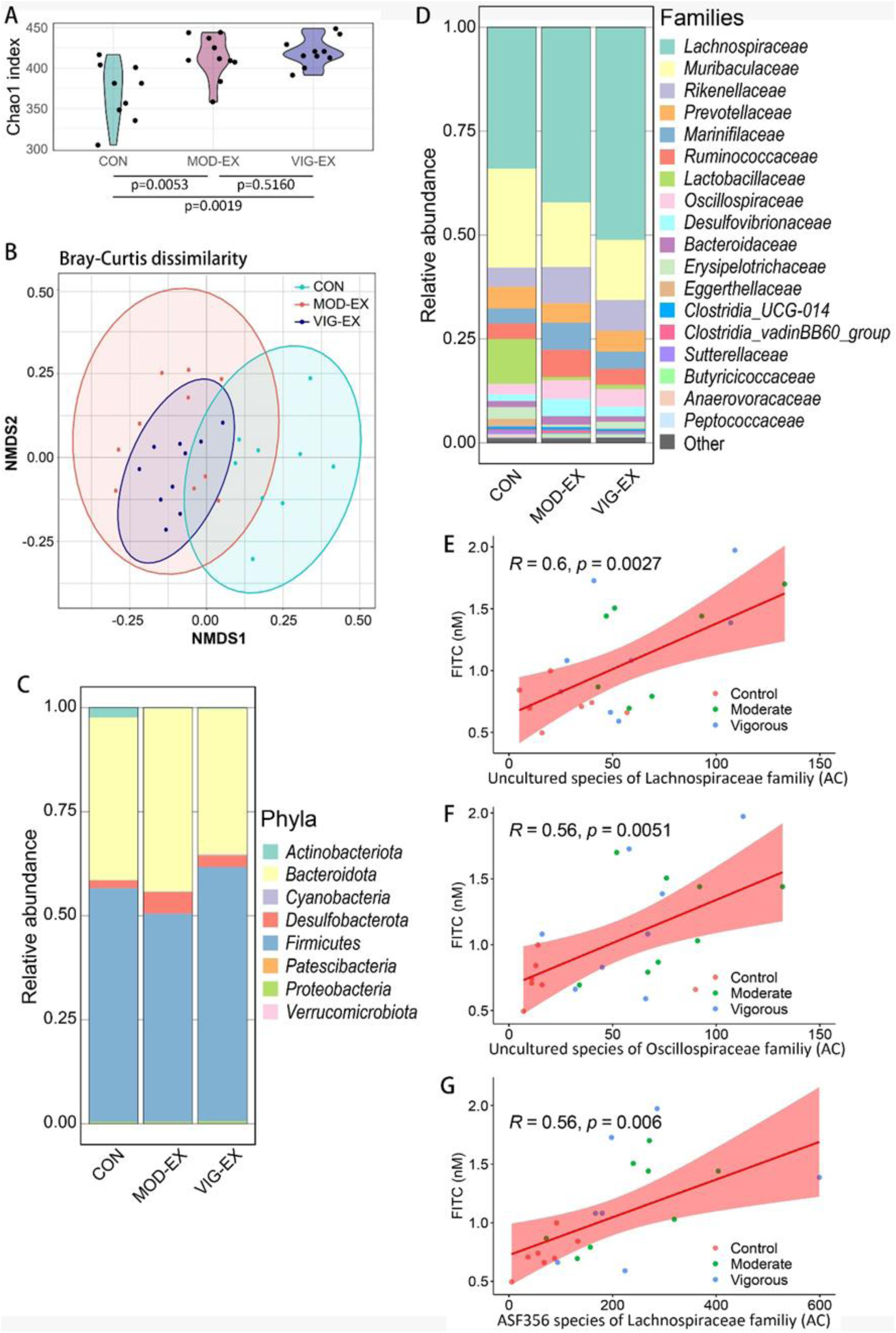
The effect of exercise training on caecal microbiota diversity and composition. (**A**) Alpha diversity depicted as Chao1 index. (**B**) Beta diversity depicted as NMDS plot based on Bray–Curtis dissimilarity matrix. Each point represents a single sample and the closed areas represent confidence ellipses. The statistical significance among the groups is determined with permutational multivariate analysis of variance (PERMANOVA), adjusted for exercise status. (**C**, **D**) Bacterial taxonomic composition in the mouse cecum following 6 weeks of exercise training at phylum and family levels. The relative abundances are displayed as mean of samples. “Other” in (D) includes all the families with less than 0.2% abundance and unidentified families. (**E**, **F**, **G**) Pearson’s correlation analyses between the specific gut microorganism abundance and serum FITC-dextran concentration of CON, MOD-EX and VIG-EX. n = 9 (CON) or 10 (MOD-EX and VIG-EX). CON: Control; MOD-EX: moderate exercise; VIG-EX: vigorous exercise; NMDS: non-metric multi-dimensional scaling; AC : abundance counts.

The microbiome of all groups was dominated by bacteria belonging to the phylae *Bacteroidota* or *Firmicutes,* and MOD-EX and VIG-EX only caused a decrease of *Actinobacteriota* (from 2.31 ± 0.90% to 0.02 ± 0.00% and 0.03 ± 0.01%, respectively) (Fig. 5C). However, the microbial composition of MOD-EX and VIG-EX differed from that of CON at family level. *Lactobacillaceae* accounted for 10.3 ± 4.0% of total caecal bacteria in CON but their abundance was reduced to less than 1% in MOD-EX and VIG-EX (Fig. 5D).

Microbiome taxonomic abundance was further analysed using LEfSe analyses which identifies discriminating bacterial taxa between groups based on statistical significance and biological relevance. Results were ranked by their linear discriminant analysis (LDA) score. Also here, the abundance of *Lactobacillaceae* (of the phylum *Firmicutes*) was identified as the main distinction in CON, whereas in MOD-EX and VIG-EX the strongest associations were related to *Rikenellaceae* (of the phylum *Bacteroidota*) and *Lachnospiraceae* (of the phylum *Firmicutes*), respectively (Supplementary Fig. S8). When analysing the taxa below family level, *Prevotellaceae UCG-001* from the phylum *Bacteroidota* was identified as the most enriched genus in MOD-EX, followed by *Rikenellaceae RC9* (Supplementary Fig. S9, S10). In VIG-EX, *Lachnospiraceae UCG-001* and the family members *Roseburia*, *ASF356* and *Acetatifactor* were identified as the most enriched genera (Supplementary Fig. S9, S10). Pearson’s correlation analyses were carried out between the relative microbiota abundance and the intestinal permeability measured with FITC-Dextran. Three species from the family of *Lachnospiraceae* and the genus *Butyricicoccus* were found to positively correlate with the serum FITC-dextran concentration (p ≤ 0.01, Fig. 5E-G), indicating a link between these species and intestinal permeability.

## 4. Discussion

Strenuous exercise can dysregulate the intestinal homeostasis that is tightly linked to the function of the intestinal barrier, affecting nutrient absorption and the defence against pathogens and toxins [20]. Here we investigated the intestinal alterations induced by exercise training of different intensities (moderate and vigorous) and outlined the mechanisms behind exercise-induced intestinal injury. This study found that continuous moderate intensity exercise imposed greater damage on the mouse intestine than vigorous intensity exercise with intermittent rest days. Increased intestinal SCFA production observed with vigorous exercise training may hereby have compensated the exercise-induced intestinal damage.

In this study, the difference in exercise intensity was reflected in different average running speeds, durations and frequencies. The average running speed of VIG-EX was 43-49% higher than that of MOD-EX but VIG-EX only ran 30% of the MOD-EX running time. This resulted in a lower total distance run by VIG-EX compared to MOD- EX (11,835 m vs 26,174 m). Notably, 4 weeks of MOD-EX (16,674 m) was sufficient to increase intestinal permeability, which was first observed at week 6 in VIG-EX (11,835 m). Substantial differences in running speed and distance may lead to distinct changes in energy metabolism, which can be the underlying cause of the differences observed in intestinal parameters (e.g. microbiota composition and metabolite levels) between the two exercise groups. Exercise training can modulate energy metabolism by affecting caloric intake and our recent study indeed demonstrated that mice that trained at a moderate treadmill running speed ate more high fat high cholesterol diet than untrained mice [17]. Strenuous exercise in humans, on the other hand, suppresses appetite and food intake [21]. Here, no significant differences in body weight or food intake between the groups were observed. As we did not assess the feeding events throughout training and rest days separately, we cannot exclude that different exercise training intensities affected dietary patterns.

As a response to heat stress during exercise, heart rate and blood flow are increased to accelerate heat dissipation. Molecular chaperones, such as heat shock proteins HSP-70 and HSP-90, dissociate from HSF-1 to perform their reparative roles by assisting protein refolding and by promoting the degradation of misfolded proteins [22]. This also allows HSF-1 to stimulate the transcription of target genes that help to cope with such proteotoxic stress [23]. We observed that the expression of HSP-70 and HSF-1 in the colon was elevated one hour after the final training, indicating that the effect of heat stress caused by both moderate and vigorous exercise continues after exercising and may cause (intestinal) tissue damage. Interestingly, after 6 weeks, only MOD-EX elevated the level of serum CRP, potentially suggesting more severe intestinal damage and systemic inflammation following moderate rather than vigorous exercise training. This was in line with elevated morphological scores and significantly shortened villi with MOD-EX. However, circulating pro-inflammatory cytokines and chemokines were undetectable or not significantly elevated after 6 weeks, indicating overall low systemic inflammation in these healthy mice. It may be of interest to study the inflammatory response to moderate and vigorous exercise over time to establish whether CRP levels in MOD-EX subside after an initial induction or whether VIG-EX also elevates circulating CRP after a longer training period. Here, it would also be meaningful to assess whether the frequency of intermittent rest days can modulate whole-body and intestinal inflammation and whether these effects are directly linked to the intestinal barrier function. A clinical study showed a link between exercise intensity and small intestinal permeability, however, all groups trained at the same frequency, making it difficult to assess the importance of rest days in the recovery of the intestine [12]. Of note, high-intensity interval training was found to have superior beneficial effects on cardiovascular function and the metabolic syndrome compared to moderate intensity continuous training [71,72]. Nonetheless, clinical studies on the effects of exercise intensity and frequency on intestinal injury and pro-inflammatory disease drivers are lacking.

Functionally, both MOD-EX and VIG-EX after 6 weeks increased intestinal leakage which correlated positively with rectal temperature. This increase in intestinal permeability is likely causally linked to impaired intestinal epithelial tight junction expression and localization. After 6 weeks of exercise training that induced elevated rectal temperatures, occludin and ZO-1 protein expression was lower in MOD-EX colon compared to CON, implicating heat stress as the causal factor. Another *in vivo* study also showed that heat stress can induce severe intestinal barrier damage [24]. Interestingly, claudin-3 protein expression increased after 6 weeks of exercise. This might be a compensatory mechanism to help maintain the integrity of the tight junctions, since Poritz *et al.* demonstrated that the increase of claudin-1 compensated the loss of ZO-1 in a murine colitis model [25]. *In vitro* we previously observed an increase of AJ protein E-cadherin, accompanied by a decrease of the TJ proteins ZO-1, claudin-3 and occludin, in a hypoxia- and heat-exposed Caco-2/HT-29 co-culture model [13]. Conversely, several other studies described an exercise-induced upregulation of TJ proteins (e.g. claudin-1, claudin-5 and occludin) as markers of enhanced barrier function in the gut and even in other organs with mild exercise [26]. Together, these studies highlight the complexity of the intestinal tight junction network in response to physical exercise.

Both intensities of exercise training altered the the gut microbiota composition which may directly affect the integrity of the intestinal epithelium. For example, the effects of *Lactobacillus* and its derivatives on preserving TJ protein expression and distribution in the intestinal tract have been proven in various *in vivo* and *ex vivo* studies [27–29]. Here, both intensities of exercise training diminished the presence of caecal *Lactobacillus*, potentially contributing to the observed intestinal barrier disruption. In future studies, *Lactobacillus*-enriched diets or gut microbiota transplants could be introduced to observe whether the addition of *Lactobacillus* can counteract the intestinal leakage from moderate to vigorous exercise training.

SCFAs are the products of microbial fermentation of non-digestible oligosaccharides [30,31]. These metabolites, like butyrate, can improve gut barrier function by enhancing mucin and TJ protein ZO-1, occludin and claudin-1 expression [32,33], and exert a general protective effect on the host by enhancing immunity and their anti-inflammatory capacity under stress [34–37]. Bacteria such as *Lachnospiraceae*, *Prevotellaceae*, and *Butyricicoccaceae*, which are strongly associated with SCFA production [38], were significantly more abundant in both exercise groups. This may arise from the decrease of commensal lactobacilli, lifting the inhibitory effect of environmental acidification on the growth of e.g. *Lachnospiraceae* [39]. Moreover, the increased activity of endogenous antioxidants after SCFA supplementation has been demonstrated in *in vitro* and *in vivo* studies [40,41], suggesting that SCFAs also modulate the oxidative stress pathways. The increase of SCFA production in the gut may also contribute to increased muscle strength/growth and whole-body energy expenditure through increased fatty acid oxidation [42–44], or interact specifically with GPR41 and GPR43, stimulating the secretion of incretins involved in whole-body metabolism [45,46]. This underlines the important reciprocal link between SCFAs and whole-body homeostasis. In fact, VIG-EX, with increased abundance of *Lachnospiraceae* spp. and significantly elevated SCFAs, which correlated with GPR41 and 43 expression levels [47], demonstrated less severe intestinal damage than MOD-EX. Accordingly, we speculate that exercise training with intermittent rest days (VIG-EX) is more favourable (or less harmful) to the colonization and reproduction of these SCFA-producing species than daily exercise (MOD-EX). While exercise can cause intestinal epithelial damage and activation of HSPs, changes to the microbiota composition and the enhanced production of SCFAs may be part of the positive adaptation to chronic exercise training that compensate for exercise-induced GI stress.

In addition, 6 weeks of VIG-EX significantly reduced caecal amino acid concentrations. This decrease could be related to increased intestinal leakiness or associated with elevated SCFA production, since total caecal SCFAs correlated negatively with total caecal amino acids. Amino acids can be utilized by intestinal microorganisms to synthesize structural proteins and serve as precursors of SCFAs [48]. Increased leakiness, on the other hand, could also promote increased influx of muscle-derived lactate during exercise that can also serve as precursor of microbially produced SCFAs, thus providing extra energy by subsequent fatty acid oxidation [49]. As no significantly increased caecal lactate levels were observed after exercise, it can be speculated that lactate was quickly converted and/or metabolized soon after exercise. Lactate production increases significantly with increasing running speed in mice [50]. In line, we speculate that in VIG-EX, higher circulating lactate levels during and immediately after the vigorous exercise contributed to the increased SCFA production. This and other exercise-induced fluxes of metabolites and their effect on gut microbes should be investigated further.

In summary (Fig. 6), exercise elevated the core body temperature, leading to heat stress and increased intestinal epithelial permeability characterized by disruption of the TJ network. Meanwhile, exercise training also altered the gut microbiota and reduced the abundance of *Lactobacillaceae* that may have contributed to the deteriorated intestinal status, indicating a novel cause of intestinal injury from exercise training. On the other hand, exercise training increased the abundance of SCFA-producing bacteria such as *Lachnospiraceae*, potentially as a compensatory response to alleviate intestinal injury.

**Figure 6.**
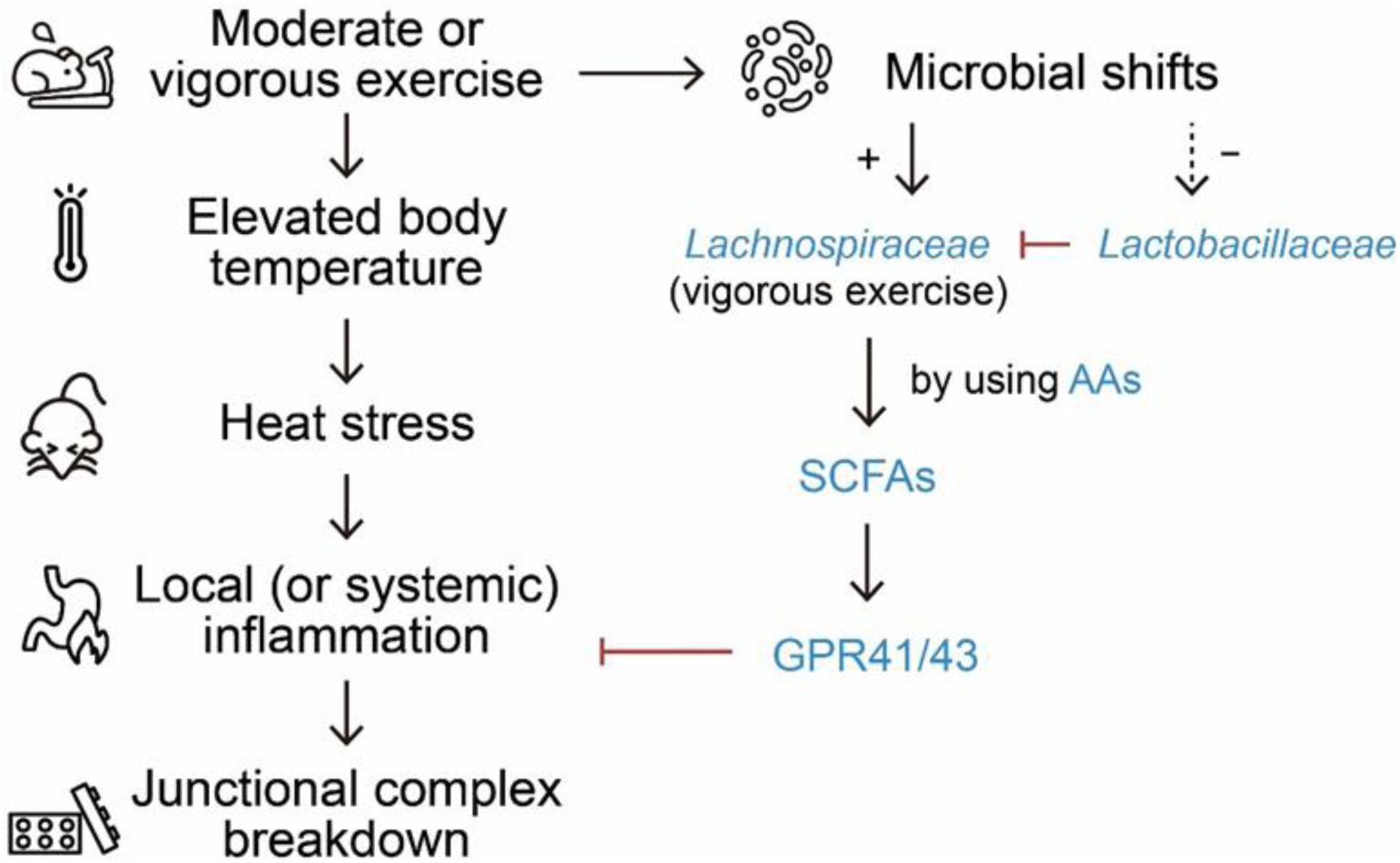
Postulated mechanism underlying the systemic and local changes in the gastrointestinal tract induced by strenuous exercise. Solid or dashed arrow: induce, produce, activate or lead to; +/-: increase/decrease; bar-headed arrow: suppress. SCFAs: short-chain fatty acids; AAs: amino acids; GPR: G-protein coupled receptor.

## Acknowledgements

We wish to gratefully acknowledge the technical and intellectual support provided by our colleagues (in alphabetical order by last names) Ingrid van Ark, Tiago Cardoso, Mara Diks, Gemma Dingjan, Thea Leusink-Muis, Lucía Peralta, Elena Sendino, Negisa Seyedtoutounchi, Koen Westphal, Yi Yang, Yuanpeng Zheng and Marit Zuurveld at Utrecht Institute of Pharmaceutical Sciences (UIPS), Utrecht University, The Netherlands.

## Author contributions

P.L., G.F., and S.B. conceived and designed research; P.L., S.K., P.C.N.R., and M.S. drafted the animal study protocol; P.L., A.K., L.W., A.C.M.P., A.V., M.G., S.T., B.M.T., and M.S. performed experiments, analysed data and interpreted results of experiments; P.L. and A.K. prepared figures; P.L. drafted manuscript; H.T., H.J.W., P.A.J.H., G.F., P.C.N.R., M.S. and S.B. edited and revised manuscript; all authors commented on and approved the final version of manuscript.

## Institutional Review Board Statement

The animal study protocol was approved by the National Committee for Animal Experiments and by the Ethics Committee on Animal Care and Experimentation of Leiden University Medical Centre (IvD Leiden). All animal experiments were performed in accordance with the Institute for Laboratory Animal Research Guide for the Care and Use of Laboratory.

## Data Availability Statement

The datasets used and/or analysed during the current study are available from the corresponding author on reasonable request.

## References

1. Ruegsegger, G.N.; Booth, F.W. Health Benefits of Exercise. Cold Spring Harb Perspect Med 2018, 8, doi:10.1101/CSHPERSPECT.A029694.

2. JanssenDuijghuijsen, L.M.; Mensink, M.; Lenaerts, K.; Fiedorowicz, E.; van Dartel, D.A.M.; Mes, J.J.; Luiking, Y.C.; Keijer, J.; Wichers, H.J.; Witkamp, R.F.;, et al. The Effect of Endurance Exercise on Intestinal Integrity in Well-Trained Healthy Men. Physiol Rep 2016, 4, doi:10.14814/PHY2.12994.

3. de Oliveira, E.P.; Burini, R.C.; Jeukendrup, A. Gastrointestinal Complaints During Exercise: Prevalence, Etiology, and Nutritional Recommendations. Sports Med 2014, 44, 79, doi:10.1007/S40279-014-0153-2.

4. Ribeiro, F.M.; Petriz, B.; Marques, G.; Kamilla, L.H.; Franco, O.L. Is There an Exercise-Intensity Threshold Capable of Avoiding the Leaky Gut? Front Nutr 2021, 8, 627289, doi:10.3389/FNUT.2021.627289.

5. Costa, R.J.S.; Snipe, R.M.J.; Kitic, C.M.; Gibson, P.R. Systematic Review: Exercise-Induced Gastrointestinal Syndrome—Implications for Health and Intestinal Disease. Aliment Pharmacol Ther 2017, 46, 246–265, doi:10.1111/APT.14157.

6. Qamar, M.I.; Read, A.E. Effects of Exercise on Mesenteric Blood Flow in Man. Gut 1987, 28, 583–587, doi:10.1136/GUT.28.5.583.

7. Perko, M.J.; Nielsen, H.B.; Skak, C.; Clemmesen, J.O.; Schroeder, T. v.; Secher, N.H. Mesenteric, Coeliac and Splanchnic Blood Flow in Humans during Exercise. J Physiol 1998, 513, 907, doi:10.1111/J.1469-7793.1998.907BA.X.

8. Keirns, B.H.; Koemel, N.A.; Sciarrillo, C.M.; Anderson, K.L.; Emerson, S.R. Exercise and Intestinal Permeability: Another Form of Exercise-Induced Hormesis? Am J Physiol Gastrointest Liver Physiol 2020, 319, G512–G518, doi:10.1152/AJPGI.00232.2020/ASSET/IMAGES/LARGE/ZH30102078410001.JPEG.

9. Ortega, A.D.S.V.; Szabó, C. Adverse Effects of Heat Stress on the Intestinal Integrity and Function of Pigs and the Mitigation Capacity of Dietary Antioxidants: A Review. Animals 2021, Vol. 11, Page 1135 2021, 11, 1135, doi:10.3390/ANI11041135.

10. Santos, R.R.; Awati, A.; Roubos-van den Hil, P.J.; Tersteeg-Zijderveld, M.H.G.; Koolmees, P.A.; Fink-Gremmels, J. Quantitative Histo-Morphometric Analysis of Heat-Stress-Related Damage in the Small Intestines of Broiler Chickens. Avian Pathology 2015, 44, 19–22, doi:10.1080/03079457.2014.988122/SUPPL_FILE/CAVP_A_988122_SM6225.DOCX.

11. Yeh, Y.J.; Law, L.Y.L.; Lim, C.L. Gastrointestinal Response and Endotoxemia during Intense Exercise in Hot and Cool Environments. Eur J Appl Physiol 2013, 113, 1575–1583, doi:10.1007/S00421-013-2587-X/FIGURES/4.

12. Pals, K.L.; Chang, R.T.; Ryan, A.J.; Gisolfi, C. v. Effect of Running Intensity on Intestinal Permeability. J Appl Physiol 1997, 82, 571–576, doi:10.1152/JAPPL.1997.82.2.571/ASSET/IMAGES/LARGE/JAPP0523305.JPEG.

13. Lian, P.; Braber, S.; Varasteh, S.; Wichers, H.J.; Folkerts, G. Hypoxia and Heat Stress Affect Epithelial Integrity in a Caco-2/HT-29 Co-Culture. Sci Rep 2021, 11, doi:10.1038/S41598-021-92574-5.

14. Radons, J. The Human HSP70 Family of Chaperones: Where Do We Stand? Cell Stress Chaperones 2016, 21, 379, doi:10.1007/S12192-016-0676-6.

15. Fehrenbach, E.; Niess, A.M.; Schlotz, E.; Passek, F.; Dickhuth, H.H.; Northoff, H. Transcriptional and Translational Regulation of Heat Shock Proteins in Leukocytes of Endurance Runners. J Appl Physiol (1985) 2000, 89, 704–710, doi:10.1152/JAPPL.2000.89.2.704.

16. Fehrenbach, E.; Passek, F.; Niess, A.M.; Pohla, H.; Weinstock, C.; Dickhuth, H.H.; Northoff, H. HSP Expression in Human Leukocytes Is Modulated by Endurance Exercise. Med Sci Sports Exerc 2000, 32, 592–600, doi:10.1097/00005768-200003000-00007.

17. Schönke, M.; Ying, Z.; Kovynev, A.; In het Panhuis, W.; Binnendijk, A.; van der Poel, S.; Pronk, A.C.M.; Streefland, T.C.M.; Hoekstra, M.; Kooijman, S.; et al. Time to Run: Late Rather than Early Exercise Training in Mice Remodels the Gut Microbiome and Reduces Atherosclerosis Development. FASEB J 2023, 37, doi:10.1096/FJ.202201304R.

18. Monda, V.; Villano, I.; Messina, A.; Valenzano, A.; Esposito, T.; Moscatelli, F.; Viggiano, A.; Cibelli, G.; Chieffi, S.; Monda, M.;, et al. Exercise Modifies the Gut Microbiota with Positive Health Effects. Oxid Med Cell Longev 2017, 2017, doi:10.1155/2017/3831972.

19. Helander, H.F.; Fändriks, L. Surface Area of the Digestive Tract – Revisited. http://dx.doi.org/10.3109/00365521.2014.898326 2014, 49, 681–689, doi:10.3109/00365521.2014.898326.

20. Lian, P.; Braber, S.; Garssen, J.; Wichers, H.J.; Folkerts, G.; Fink-Gremmels, J.; Varasteh, S. Beyond Heat Stress: Intestinal Integrity Disruption and Mechanism-Based Intervention Strategies. Nutrients 2020, 12, 734.

21. Dorling, J.; Broom, D.R.; Burns, S.F.; Clayton, D.J.; Deighton, K.; James, L.J.; King, J.A.; Miyashita, M.; Thackray, A.E.; Batterham, R.L.;, et al. Acute and Chronic Effects of Exercise on Appetite, Energy Intake, and Appetite-Related Hormones: The Modulating Effect of Adiposity, Sex, and Habitual Physical Activity. Nutrients 2018, 10, doi:10.3390/NU10091140.

22. Whitley, D.; Goldberg, S.P.; Jordan, W.D. Heat Shock Proteins: A Review of the Molecular Chaperones. J Vasc Surg 1999, 29, 748–751, doi:10.1016/S0741-5214(99)70329-0.

23. Zhang, Y.; Chou, S.D.; Murshid, A.; Prince, T.L.; Schreiner, S.; Stevenson, M.A.; Calderwood, S.K. The Role Of Heat Shock Factors In Stress-Induced Transcription. Methods Mol Biol 2011, 787, 21, doi:10.1007/978-1-61779-295-3_2.

24. Koch, F.; Thom, U.; Albrecht, E.; Weikard, R.; Nolte, W.; Kuhla, B.; Kuehn, C. Heat Stress Directly Impairs Gut Integrity and Recruits Distinct Immune Cell Populations into the Bovine Intestine. Proc Natl Acad Sci U S A 2019, 116, 10333–10338, doi:10.1073/PNAS.1820130116/SUPPL_FILE/PNAS.1820130116.SD05.XLSX.

25. Poritz, L.S.; Garver, K.I.; Green, C.; Fitzpatrick, L.; Ruggiero, F.; Koltun, W.A. Loss of the Tight Junction Protein ZO-1 in Dextran Sulfate Sodium Induced Colitis. Journal of Surgical Research 2007, 140, 12–19, doi:10.1016/J.JSS.2006.07.050.

26. Shin, H.E.; Kwak, S.E.; Zhang, D. di; Lee, J.; Yoon, K.J.; Cho, H.S.; Moon, H.Y.; Song, W. Effects of Treadmill Exercise on the Regulation of Tight Junction Proteins in Aged Mice. Exp Gerontol 2020, 141, 111077, doi:10.1016/J.EXGER.2020.111077.

27. Liu, J.; Li, R.; Liu, P.; Ruiz, J.; Qin, D.; Ma, Y.; Wang, Y.; Hou, X.; Yu, L. Contribution of Lactobacilli on Intestinal Mucosal Barrier and Diseases: Perspectives and Challenges of Lactobacillus Casei. Life 2022, Vol. 12, Page 1910 2022, 12, 1910, doi:10.3390/LIFE12111910.

28. Blackwood, B.P.; Yuan, C.Y.; Wood, D.R.; Nicolas, J.D.; Grothaus, J.S.; Hunter, C.J. Probiotic Lactobacillus Species Strengthen Intestinal Barrier Function and Tight Junction Integrity in Experimental Necrotizing Enterocolitis. J Probiotics Health 2017, 5, doi:10.4172/2329-8901.1000159.

29. Samak, G.; Rao, R.; Rao, R. Lactobacillus Casei and Epidermal Growth Factor Prevent Osmotic Stress-Induced Tight Junction Disruption in Caco-2 Cell Monolayers. Cells 2021, Vol. 10, Page 3578 2021, 10, 3578, doi:10.3390/CELLS10123578.

30. Abuja, P.M.; Vinelli, V.; Biscotti, P.; Martini, D.; del Bo’, C.; Marino, M.; Meroño, T.; Nikoloudaki, O.; Calabrese, F.M.; Turroni, S.;, et al. Effects of Dietary Fibers on Short-Chain Fatty Acids and Gut Microbiota Composition in Healthy Adults: A Systematic Review. Nutrients 2022, Vol. 14, Page 2559 2022, 14, 2559, doi:10.3390/NU14132559.

31. Xu, T.; Wu, X.; Liu, J.; Sun, J.; Wang, X.; Fan, G.; Meng, X.; Zhang, J.; Zhang, Y. The Regulatory Roles of Dietary Fibers on Host Health via Gut Microbiota-Derived Short Chain Fatty Acids. Curr Opin Pharmacol 2022, 62, 36–42, doi:10.1016/J.COPH.2021.11.001.

32. Ortiz-Alvarez, L.; Xu, H.; Martinez-Tellez, B. Influence of Exercise on the Human Gut Microbiota of Healthy Adults: A Systematic Review. Clin Transl Gastroenterol 2020, 11, e00126, doi:10.14309/CTG.0000000000000126.

33. Ghosh, S.; Whitley, C.S.; Haribabu, B.; Jala, V.R. Regulation of Intestinal Barrier Function by Microbial Metabolites. Cell Mol Gastroenterol Hepatol 2021, 11, 1463, doi:10.1016/J.JCMGH.2021.02.007.

34. Hawley, J.A. Microbiota and Muscle Highway — Two Way Traffic. Nature Reviews Endocrinology 2019 16:2 2019, 16, 71–72, doi:10.1038/s41574-019-0291-6.

35. Bongiovanni, T.; Yin, M.O.L.; Heaney, L.M. The Athlete and Gut Microbiome: Short-Chain Fatty Acids as Potential Ergogenic Aids for Exercise and Training. Int J Sports Med 2021, 42, 1143–1158, doi:10.1055/A-1524-2095.

36. Antonini, M.; Conte, M. lo; Sorini, C.; Falcone, M. How the Interplay between the Commensal Microbiota, Gut Barrier Integrity, and Mucosal Immunity Regulates Brain Autoimmunity. Front Immunol 2019, 10, 1937, doi:10.3389/FIMMU.2019.01937/BIBTEX.

37. Cai, Y.; Folkerts, J.; Folkerts, G.; Maurer, M.; Braber, S. Microbiota-dependent and -independent Effects of Dietary Fiber on Human Health. Br J Pharmacol 2019, bph.14871, doi:10.1111/bph.14871.

38. Xu, Y.; Zhu, Y.; Li, X.; Sun, B. Dynamic Balancing of Intestinal Short-Chain Fatty Acids: The Crucial Role of Bacterial Metabolism. Trends Food Sci Technol 2020, 100, 118–130, doi:10.1016/J.TIFS.2020.02.026.

39. Brownlie, E.J.E.; Chaharlangi, D.; Wong, E.O.Y.; Kim, D.; Navarre, W.W. Acids Produced by Lactobacilli Inhibit the Growth of Commensal Lachnospiraceae and S24-7 Bacteria. Gut Microbes 2022, 14, doi:10.1080/19490976.2022.2046452/SUPPL_FILE/KGMI_A_2046452_SM4942.ZIP.

40. Hamer, H.M.; Jonkers, D.M.A.E.; Bast, A.; Vanhoutvin, S.A.L.W.; Fischer, M.A.J.G.; Kodde, A.; Troost, F.J.; Venema, K.; Brummer, R.J.M. Butyrate Modulates Oxidative Stress in the Colonic Mucosa of Healthy Humans. Clinical Nutrition 2009, 28, 88–93, doi:10.1016/j.clnu.2008.11.002.

41. Nielsen, D.S.G.; Jensen, B.B.; Theil, P.K.; Nielsen, T.S.; Knudsen, K.E.B.; Purup, S. Effect of Butyrate and Fermentation Products on Epithelial Integrity in a Mucus-Secreting Human Colon Cell Line. J Funct Foods 2018, 40, 9–17, doi:10.1016/J.JFF.2017.10.023.

42. Gizard, F.; Fernandez, A.; de Vadder, F. Interactions between Gut Microbiota and Skeletal Muscle. Nutr Metab Insights 2020, 13, doi:10.1177/1178638820980490/ASSET/IMAGES/LARGE/10.1177_1178638820980490-FIG1.JPEG.

43. Maruta, H.; Yamashita, H. Acetic Acid Stimulates G-Protein-Coupled Receptor GPR43 and Induces Intracellular Calcium Influx in L6 Myotube Cells. PLoS One 2020, 15, e0239428, doi:10.1371/JOURNAL.PONE.0239428.

44. Zhang, B.; Liu, H.; Liu, M.; Yue, Z.; Liu, L.; Fuchang, L. Exogenous Butyrate Regulates Lipid Metabolism through GPR41-ERK-AMPK Pathway in Rabbits. https://doi-org.proxy.library.uu.nl/10.1080/1828051X.2022.2049985 2022, 21, 473–487, doi:10.1080/1828051X.2022.2049985.

45. Christiansen, C.B.; Gabe, M.B.N.; Svendsen, B.; Dragsted, L.O.; Rosenkilde, M.M.; Holst, J.J. The Impact of Short-Chain Fatty Acids on Glp-1 and Pyy Secretion from the Isolated Perfused Rat Colon. Am J Physiol Gastrointest Liver Physiol 2018, 315, G53–G65, doi:10.1152/AJPGI.00346.2017/ASSET/IMAGES/LARGE/ZH30041874330006.JPEG.

46. Grasset, E.; Puel, A.; Charpentier, J.; Collet, X.; Christensen, J.E.; Tercé, F.; Burcelin, R. A Specific Gut Microbiota Dysbiosis of Type 2 Diabetic Mice Induces GLP-1 Resistance through an Enteric NO-Dependent and Gut-Brain Axis Mechanism. Cell Metab 2017, 25, 1075–1090.e5, doi:10.1016/J.CMET.2017.04.013.

47. Kim, M.H.; Kang, S.G.; Park, J.H.; Yanagisawa, M.; Kim, C.H. Short-Chain Fatty Acids Activate GPR41 and GPR43 on Intestinal Epithelial Cells to Promote Inflammatory Responses in Mice. Gastroenterology 2013, 145, 396–406.e10, doi:10.1053/J.GASTRO.2013.04.056.

48. Lin, R.; Liu, W.; Piao, M.; Zhu, H. A Review of the Relationship between the Gut Microbiota and Amino Acid Metabolism. Amino Acids 2017, 49, 2083–2090, doi:10.1007/S00726-017-2493-3/FIGURES/1.

49. Scheiman, J.; Luber, J.M.; Chavkin, T.A.; MacDonald, T.; Tung, A.; Pham, L.D.; Wibowo, M.C.; Wurth, R.C.; Punthambaker, S.; Tierney, B.T.;, et al. Meta-Omics Analysis of Elite Athletes Identifies a Performance-Enhancing Microbe That Functions via Lactate Metabolism. Nature Medicine 2019 25:7 2019, 25, 1104–1109, doi:10.1038/s41591-019-0485-4.

50. Odriozola, C.P.; Cordero, J.Á.G.; Daura, J.; Sanz, M.; Martínez-Blanes, J.M.; Avilés, M.Á. Reliability of Blood Lactate as a Measure of Exercise Intensity in Different Strains of Mice during Forced Treadmill Running. PLoS One 2019, 14, e0215584, doi:10.1371/JOURNAL.PONE.0215584.

